# GinsengMind: An evidence-grounded AI research agent for integrative molecular research in *Panax ginseng*

**DOI:** 10.64898/2026.11.22.768453

**Authors:** Yuru Tong, Rong Zhang, Chuanzhi Kang, Zeqian Chang, Kang Chen

## Abstract

*Panax ginseng* C. A. Meyer is one of the most extensively studied medicinal plants, with substantial pharmaceutical and economic importance owing to its diverse bioactive constituents, particularly ginsenosides. Despite considerable advances in ginseng molecular research, the fragmentation of genomic, transcriptomic, protein structural, and biochemical information remains a major challenge for investigating gene function and specialized metabolism. Here, we developed GinsengMind (https://ginsengai.nrc.ac.cn), an evidence-grounded AI research agent that integrates species-specific molecular knowledge with large language model (LLM)-guided tool orchestration and structured retrieval-augmented generation. Built upon the telomere-to-telomere reference genome of *P. ginseng*, GinsengMind integrates 77,266 genes, 154,754 transcripts, and 77,266 protein records. Comprehensive functional annotation using Swiss-Prot, eggNOG, and InterProScan covered 97.00% of the 76,947 quality-controlled proteins. The platform further incorporates 71,046 ESMFold-predicted protein structures, 7,263,004 gene expression records across 94 RNA-seq samples, and 4,813 bibliographic records. Six specialized research tools enable autonomous evidence retrieval and source-linked scientific interpretation through natural-language interaction. To evaluate its performance, we established GinsengBench, comprising 50 curated questions across six research categories. In a blinded evaluation by three experts, GinsengMind achieved a mean weighted composite score of 96.18 across ten core questions, substantially exceeding GLM-5.3 without database access (23.24; mean paired difference, 72.94; 95% confidence interval, 65.19-79.87). Across five shared questions, GinsengMind outperformed nine general-purpose LLMs on average and achieved a score (96.56) close to that of GLM-5.3 supplied with predefined evidence packages (96.88), highlighting the importance of domain-specific knowledge and autonomous evidence retrieval. A case study of the ginsenoside biosynthetic enzyme PgUGT94Q2 further illustrates how integrated molecular evidence can reveal unresolved biological relationships and identify priorities for subsequent experimental investigation. Beyond conventional database querying, GinsengMind provides a foundation for exploring potential biological associations, prioritizing candidate biosynthetic genes, and formulating experimentally testable hypotheses. We anticipate that GinsengMind will facilitate research on medicinal plant specialized metabolism, natural product biosynthesis, enzyme discovery and engineering, and synthetic biology, while offering an adaptable framework for developing domain-specialized AI research agents across other medicinal plants and scientific disciplines.

## Introduction

*Panax ginseng* C. A. Meyer is one of the most extensively studied medicinal plants, with a long history of traditional use and substantial scientific, pharmaceutical, and economic importance (1, 2). Its medicinal value is closely associated with a diverse array of bioactive constituents, particularly ginsenosides, a major class of triterpenoid saponins characterized by considerable structural and biological diversity(3). Elucidating the biosynthetic pathways responsible for these compounds is important not only for understanding specialized metabolism in ginseng but also for identifying biosynthetic enzymes and developing alternative production strategies through synthetic biology. Advances in genome sequencing, transcriptomic analysis, and functional characterization have substantially expanded our understanding of ginsenoside biosynthesis (4, 5). In particular, the identification of oxidosqualene cyclases, cytochrome P450 monooxygenases, and UDP-dependent glycosyltransferases has provided insights into the formation, oxidation, and glycosylation of ginsenoside scaffolds (6–9). These developments have generated extensive molecular resources and created new opportunities for investigating gene function, enzyme diversity, and the molecular basis of specialized metabolism in *P. ginseng*.

The accumulation of genomic, transcriptomic, and biochemical information has stimulated the development of specialized resources for ginseng research(10). The Ginseng Genome Database and PanaxGDB have provided genomic annotations, transcriptomic information, and analytical tools for investigating *Panax* species (11, 12). More recently, the telomere-to-telomere (T2T) reference genome of *P. ginseng*, comprising 24 chromosomes and 77,266 predicted protein-coding genes, has established an improved genomic foundation for systematic molecular investigations(5). In parallel, GinDB-AI has expanded access to curated Panax-derived compounds and introduced deep learning-based approaches for predicting molecular properties and biological activities (13). Although these resources have substantially advanced ginseng research, they provide complementary information across different biological domains. Investigating a particular gene or biosynthetic enzyme therefore often requires the integration of genomic, transcriptomic, protein functional, structural, and experimental evidence from multiple sources. Establishing consistent and traceable relationships among these data types is essential for the systematic interpretation of gene function and specialized metabolism in *P. ginseng*.

Sequence similarity, orthology-based annotation, and conserved domain analysis provide complementary insights into potential protein functions(14), whereas transcriptomic profiles reveal patterns of gene expression across tissues and experimental conditions. Recent advances in protein structure prediction have further expanded the molecular evidence available for investigating protein architecture and structural features potentially associated with biological function(15). However, these sources of information differ in their biological significance and levels of certainty.

Sequence-based annotations do not necessarily establish catalytic activity, expression associations cannot independently demonstrate biochemical function, and predicted structures do not directly determine substrate specificity. Moreover, differences among genome versions, transcript isoforms, and experimentally characterized sequence identifiers can complicate the mapping of previously reported enzymes to contemporary reference genomes. Integrating these complementary data types within a unified and traceable framework therefore enables a more comprehensive assessment of candidate gene functions, facilitates the identification of potentially meaningful biological associations, and provides a foundation for formulating experimentally testable hypotheses.

Recent advances in large language models (LLMs) offer opportunities to change how researchers access and interpret complex biological information. Rather than navigating independent databases through predefined search fields, natural-language interfaces can enable researchers to formulate biological questions and explore relevant information across multiple data types. Emerging systems, including PlantGPT, DeepPGDB, PlantXBot, and CucurbitAgent, have demonstrated the potential of LLM-based approaches and AI agents for plant functional genomics and multi-omics data exploration (16–19). Nevertheless, general-purpose LLMs may generate incorrect gene identifiers, unsupported functional assignments, or unverifiable literature references, particularly when addressing species-specific molecular questions. These limitations are especially consequential in medicinal plant research, where distinguishing experimentally characterized biosynthetic reactions from computationally inferred enzyme functions is essential for reliable scientific interpretation. Integrating LLM-guided tool orchestration with structured retrieval and traceable evidence provides a promising approach to addressing these challenges. Such systems can move beyond conventional information retrieval by connecting complementary observations, highlighting unresolved relationships, and assisting researchers in formulating experimentally testable hypotheses. However, their scientific value must be established through reproducible evaluation of retrieval accuracy, evidence reliability, and performance on realistic research tasks.

Here, we developed GinsengMind (https://ginsengai.nrc.ac.cn), an evidence-grounded AI research agent for *P. ginseng*, to establish an integrated framework for molecular knowledge exploration and hypothesis-driven research in medicinal plants. Built upon the T2T reference genome, GinsengMind organizes gene, transcript, and protein information together with updated functional annotations, transcriptomic expression profiles, predicted three-dimensional protein structures, and supporting literature. A central feature of the platform is the association of complementary functional and structural evidence at the individual-protein level, enabling integrated exploration of protein sequence, annotation, expression, and predicted structure. Rather than treating conversational AI as an auxiliary database function, GinsengMind employs LLM-guided tool orchestration and structured retrieval-augmented generation as its primary mechanism for accessing and interpreting interconnected molecular information. We further introduce GinsengBench as a domain-specific evaluation framework for assessing the accuracy and traceability of AI-assisted scientific responses, and use the previously characterized glycosyltransferase PgUGT94Q2 as a representative case for examining the integration and interpretation of evidence relevant to ginsenoside biosynthesis. By connecting molecular resources with evidence-grounded scientific interaction, GinsengMind aims to support candidate biosynthetic gene prioritization, exploration of potential biological associations, and the formulation of testable hypotheses. Beyond ginseng, this work provides a framework for developing domain-specialized AI research agents with potential applications in medicinal plant specialized metabolism, natural product biosynthesis, enzyme discovery and engineering, and synthetic biology, while offering a methodological reference for other knowledge-intensive scientific disciplines.

## Results

### Architecture and evidence-grounded design of GinsengMind

GinsengMind (https://ginsengai.nrc.ac.cn) was developed as a dialogue-centered AI research platform for *Panax ginseng*, integrating a species-specific molecular knowledge base with large language model (LLM)-guided retrieval and evidence management (Fig. 1). The platform adopts a layered architecture comprising a conversational interface, an agent orchestration layer, modular research tools, and structured molecular resources. Unlike conventional database interfaces that rely primarily on independent search operations, GinsengMind uses natural-language queries as a primary entry point for accessing interconnected biological information. Dedicated interfaces for genes, proteins, expression profiles, and literature provide complementary access to the same underlying resources.

**Figure 1.**
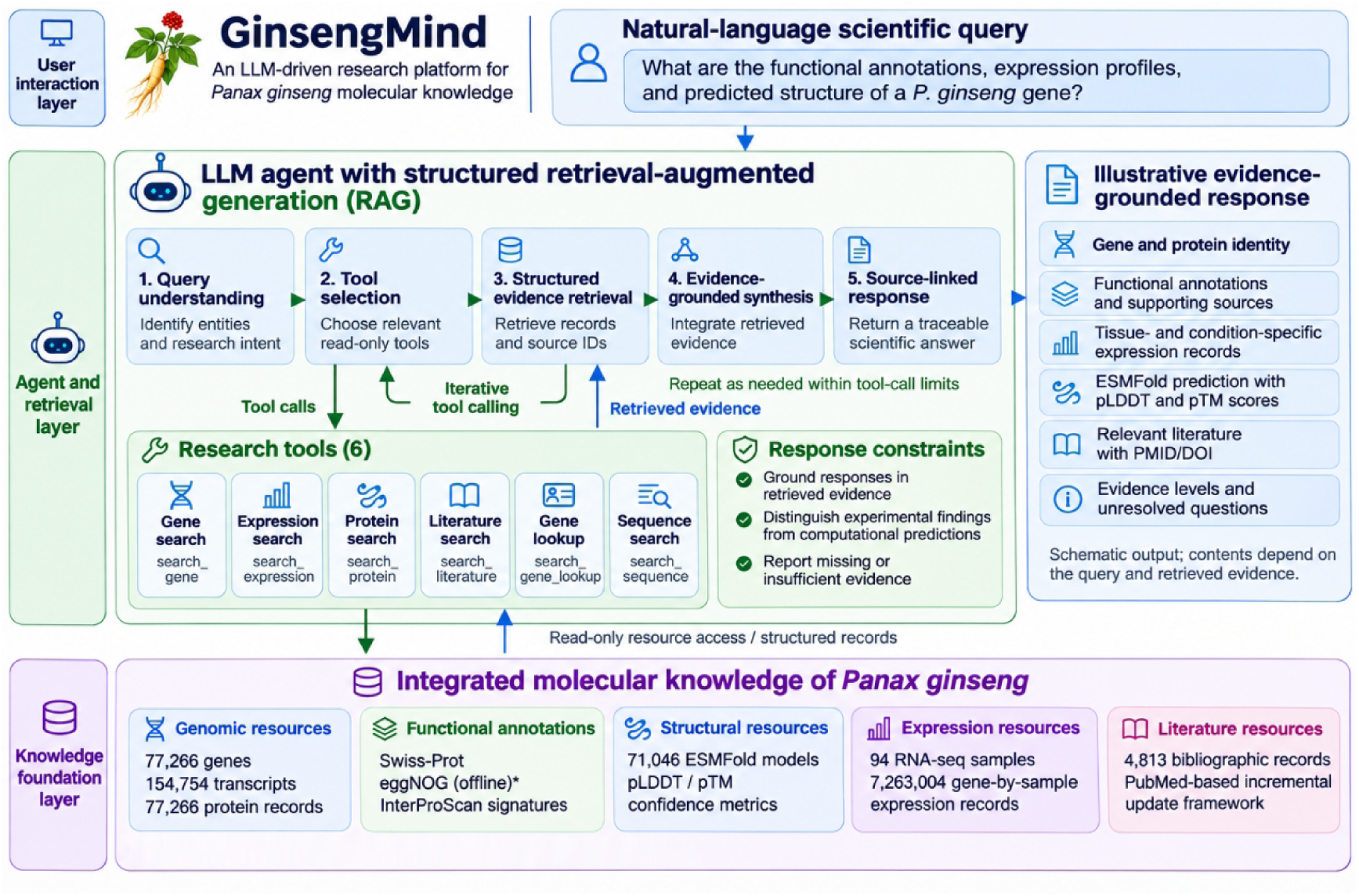
Three-layer architecture and evidence-grounded research workflow of GinsengMind.

At the core of the system, an LLM-guided agent interprets user queries and selects appropriate retrieval tools according to the requested biological information. Six research tools were implemented: search_gene, search_expression, search_protein, search_literature, search_gene_lookup, and search_sequence. These tools provide access to genomic records, expression measurements, protein annotations and structures, bibliographic information, gene identifiers, and sequence similarity searches, respectively. Tool execution is mediated through predefined interfaces and validated parameters, with database operations restricted to authorized retrieval functions. Retrieved records are subsequently organized with source identifiers and associated evidence, providing structured context for response generation. This architecture implements a form of structured retrieval-augmented generation in which scientific responses are informed by domain-specific database records rather than relying exclusively on the model’s pretrained knowledge.

Initial functional tests demonstrated the ability of the agent to retrieve and interpret reference gene information. For example, a query concerning *pg_10000006* retrieved the corresponding genomic record, including its chromosomal location, gene type, and reference annotation source. The generated response also distinguished the existence of a genomic annotation from experimentally established gene function. Additional local tests supported functional keyword retrieval and the integration of sequence similarity results with expression queries. These observations demonstrate the operational feasibility of connecting LLM-guided tool selection with structured molecular resources. The broader reliability of this approach across scientific question types is evaluated separately using GinsengBench.

GinsengMind is an LLM-driven molecular research platform for *Panax ginseng*, comprising three interconnected layers: user interaction, agent-guided retrieval, and an integrated molecular knowledge foundation. Researchers submit natural-language questions through a conversational interface, where an LLM-based agent interprets queries, selects appropriate research tools, retrieves structured molecular records, and synthesizes evidence-grounded responses. Six specialized tools support gene identification, functional annotation retrieval, sequence similarity searches, expression profiling, protein information retrieval, and literature searches. The agent employs an iterative tool-calling workflow, with retrieved information used to support response generation and evidence traceability. The underlying knowledge foundation integrates T2T reference genome annotations, protein functional annotations, ESMFold-predicted structures, transcriptomic expression profiles, and scientific literature. Evidence-management mechanisms retain source information and distinguish computational predictions from curated records while identifying missing or insufficient evidence. The response panel schematically illustrates the integration of molecular information and supporting evidence.

### An integrated molecular knowledge foundation for *Panax ginseng*

The molecular knowledge foundation of GinsengMind was constructed using the recently published telomere-to-telomere reference genome of P. ginseng. Integration of the reference annotations yielded 77,266 gene records, 154,754 transcript records, and 77,266 predicted protein records. Of the annotated transcripts, 145,357 were classified as protein-coding transcripts and 9,397 as long non-coding RNAs. Original identifiers and reference annotation information were retained to maintain correspondence between the integrated records and their genomic sources.

To establish relationships among genes, transcripts, and proteins, protein records were associated with their corresponding genes, and potential protein–transcript relationships were evaluated using coding-sequence compatibility. Among 77,266 protein records, 71,767 (92.88%) were assigned to uniquely compatible transcripts, whereas 5,491 (7.11%) had ambiguous transcript associations and eight remained unmapped. Ambiguous relationships were preserved without arbitrary assignment, allowing unresolved mappings to be distinguished from uniquely inferred associations. Protein sequence quality control further identified 319 sequences containing internal termination symbols, leaving 76,947 sequences suitable for downstream functional annotation and structure prediction.

To complement the reference genomic resources, GinsengMind incorporates a transcriptomic expression atlas comprising 94 RNA-seq samples and 7,263,004 gene-by-sample expression entries. The atlas organizes expression information together with sample metadata, including dataset origin, tissue classification, experimental conditions, and quality-control status. Expression records can be retrieved using reference gene identifiers and examined within their corresponding experimental datasets. Retaining dataset-specific metadata and quality indicators is particularly important because expression values derived from independent studies may not be directly comparable without appropriate normalization and consideration of experimental differences.

A literature resource comprising 4,813 bibliographic records was established to support evidence-based retrieval of published research on *P. ginsen*g. To maintain the currency of scientific evidence, a PubMed-based incremental synchronization framework was designed to support daily automated literature updates, including the identification of newly indexed publications, reconciliation of revised records, deduplication, and tracking of retracted or deleted publications. This framework enables the literature resource to be maintained as an evolving knowledge base for evidence-grounded scientific querying.

### Integration of protein functional annotations and predicted structures

To establish a functional and structural resource for the *P. ginseng* proteome, 76,947 quality-controlled protein sequences were subjected to complementary annotation analyses using Swiss-Prot, eggNOG-mapper, and InterProScan. A total of 74,641 proteins (97.00%) had at least one annotation record from these resources, whereas 2,306 proteins (3.00%) had no reported matches. Sequence similarity searches against Swiss-Prot identified homologous matches for 53,938 proteins (70.10%), while eggNOG-mapper provided functional annotation records for 70,044 proteins (91.03%). InterProScan identified sequence features in 73,067 proteins (94.96%), providing extensive coverage of conserved protein families, domains, and associated functional classifications.

Integration of the three annotation datasets identified 53,421 proteins (69.43% of the quality-controlled proteome) with records from all three resources. These overlapping annotations provide complementary evidence for evaluating potential protein functions, although agreement among annotation sources does not necessarily establish a specific biochemical activity. By retaining the original database accessions, functional descriptions, and annotation sources, GinsengMind enables individual functional assignments to be examined in relation to their supporting computational evidence.

To extend functional annotation beyond sequence-derived information, large-scale three-dimensional structure prediction was performed using ESMFold. A total of 71,046 predicted protein structures were generated for proteins within the computationally processed length range of up to 800 amino acids, representing approximately 91.9% of the 77,266 original protein records. Protein structure models were retained in PDB format, together with prediction confidence information, including pLDDT and pTM metrics. Structural files were transferred to the test server, where file integrity, representative three-dimensional visualization, and PDB retrieval were verified. Proteins exceeding 800 amino acids were not included in the completed prediction set, and their absence from the current structural resource was retained as a coverage limitation.

A central feature of the integrated protein resource is the association of predicted three-dimensional structures with sequence-derived functional annotations at the individual-protein level. This organization allows researchers to examine Swiss-Prot homology matches, eggNOG functional assignments, InterPro protein families and domains, and ESMFold structural models within a common molecular record. For enzymes involved in specialized metabolism, such as cytochrome P450 monooxygenases and UDP-dependent glycosyltransferases, this integration provides complementary information for assessing candidate protein functions and exploring potentially informative structural features. Nevertheless, predicted structures and sequence-derived annotations remain computational evidence and do not independently establish catalytic activity or substrate specificity.

### Benchmarking GinsengMind with GinsengBench

To systematically evaluate the performance of GinsengMind in domain-specific scientific question answering, we developed GinsengBench, a benchmark comprising 50 curated questions that cover gene information, protein annotation and structure, gene expression, literature retrieval, cross-module knowledge integration, and evidence boundaries. Each question was accompanied by reference facts and evidence-based inference boundaries to assess both the correctness of retrieved information and the validity of scientific interpretations. A core evaluation set of ten questions spanning all six categories was selected for expert assessment. A subset of five shared questions was further used for cross-model comparisons and fixed-evidence experiments. In total, 90 responses, comprising 10 from GinsengMind, 10 from No-database GLM, 45 from nine general-purpose large language models accessed through OpenRouter, and 25 from five fixed-evidence model configurations, were independently evaluated by three experts in a blinded manner. Responses were scored in four dimensions: factual accuracy, completeness, relevance, and traceability. The absolute-agreement intraclass correlation coefficient for the weighted composite scores was 0.522 for individual raters and 0.766 for the average of three raters.

Across the ten core benchmark questions, GinsengMind achieved a mean weighted composite score of 96.18, compared with 23.24 for GLM-5.3 operating without access to the GinsengMind database or retrieval tools. The mean paired difference was 72.94 points (95% question-cluster bootstrap confidence interval [CI], 65.19–79.87; Fig. 2C). Improvements were consistently observed across all four evaluation dimensions, with mean differences of 71.70, 87.37, 53.83, and 80.10 points in factual accuracy, completeness, relevance, and traceability, respectively. The largest improvement was observed in completeness, highlighting the ability of GinsengMind to retrieve and incorporate the specific information required to answer domain-specific questions. GinsengMind received higher mean composite scores for all ten questions. This advantage was consistent across the three experts and remained evident when any individual expert was excluded from the analysis. An exploratory exact paired sign test further supported the robustness of the observed difference (two-sided P = 0.00195).

**Figure 2.**
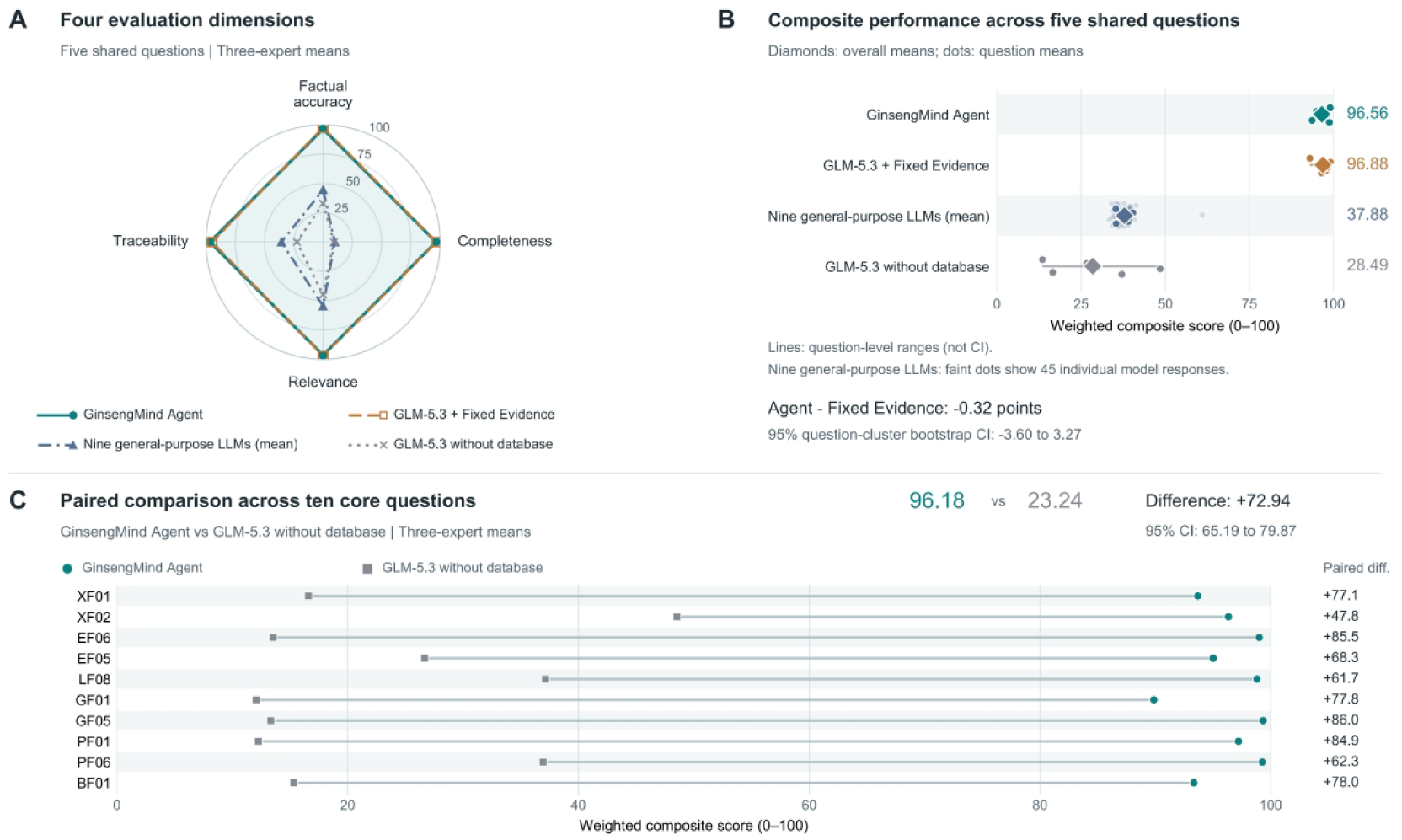
Benchmarking GinsengMind on evidence-grounded *Panax ginseng* research tasks. (A) Radar plot comparing GinsengMind Agent, nine general-purpose large language models accessed through OpenRouter (OpenRouter Nine), GLM-5.3 without database access (No-database GLM), and GLM-5.3 supplied with predefined evidence packages (Fixed Evidence B) across five shared benchmark questions. Responses were independently assessed by three experts in four dimensions: factual accuracy, completeness, relevance, and traceability, each scored on a scale of 0–100. (B) Distributions of weighted composite scores across the five shared questions. The composite score was calculated using weights of 40%, 20%, 20%, and 20% for factual accuracy, completeness, relevance, and traceability, respectively. Diamonds indicate overall means, dots represent question-level means, and horizontal lines indicate the range of question-level scores. For OpenRouter Nine, question-level scores were additionally averaged across the nine models, with faint dots representing the 45 individual model responses. (C) Paired comparison of GinsengMind Agent and No-database GLM across ten core benchmark questions. Each pair of connected dots represents the mean scores assigned by three experts to responses to the same question. Positive differences indicate higher scores for GinsengMind. The 95% confidence intervals for paired mean differences in (B) and (C) were estimated using 10,000 question-cluster bootstrap resamples, conditional on the three participating experts.

To extend the evaluation beyond the no-database GLM baseline, we compared GinsengMind with nine general-purpose large language models across five shared benchmark questions (Fig. 2A,B). GinsengMind achieved a mean weighted composite score of 96.56, whereas the corresponding scores for OpenRouter Nine and No-database GLM were 37.88 and 28.49, respectively. In particular, GinsengMind maintained consistently high mean scores in factual accuracy (96.53), completeness (96.53), relevance (96.67), and traceability (96.53). These results demonstrate that, within the evaluated question set, integration of a specialized research database with autonomous retrieval tools enabled GinsengMind to provide more complete, accurate, and traceable scientific responses than the tested general-purpose model configurations.

To further investigate the contribution of domain-specific evidence to answer quality, we conducted fixed-evidence experiments in which models received predefined evidence packages instead of retrieving information autonomously through database tools. Across the five shared questions, the five fixed-evidence configurations achieved mean weighted composite scores ranging from 85.17 to 96.88. Notably, the fixed-evidence GLM-5.3 configuration (Fixed Evidence B) achieved a mean score of 96.88, close to that of GinsengMind (96.56), with a mean paired difference of −0.32 points (95% CI, −3.60 to 3.27; Fig. 2A,B). Although this comparison does not establish statistical equivalence, it shows that GinsengMind can achieve high expert-assessed answer quality through autonomous retrieval of domain-specific information without requiring predefined, question-specific evidence packages. Together, these findings support the feasibility of integrating structured biological databases and autonomous evidence retrieval into a specialized AI agent for ginseng research. Given the limited number of evaluated questions and differences in model deployment and inference configurations, these findings should be interpreted as a system-level assessment rather than an isolated comparison of underlying language model capabilities.

### Evidence-guided investigation of PgUGT94Q2 in ginsenoside biosynthesis

To examine the application of GinsengMind to medicinal plant specialized metabolism, we investigated the potential relationship between the T2T reference locus *pg_5001060* and PgUGT94Q2, a previously characterized UDP-dependent glycosyltransferase involved in ginsenoside biosynthesis(20) (Fig. 3). This case represents a common challenge in functional genomics: connecting experimentally characterized enzymes reported in earlier studies with contemporary reference gene annotations while integrating complementary molecular evidence. A genuine GinsengMind Chat query retrieved gene, expression, and protein information associated with *pg_5001060*, together with source-linked records and explicit statements of unresolved evidence (Fig. 3A). The response identified the locus as a protein-coding gene with UGT-related annotations but did not establish its identity as the experimentally characterized PgUGT94Q2. Literature retrieval was not executed during this interaction because the configured tool-call limit was reached; therefore, the literature findings presented below were obtained through independent database searches rather than the same Chat session.

**Figure 3.**
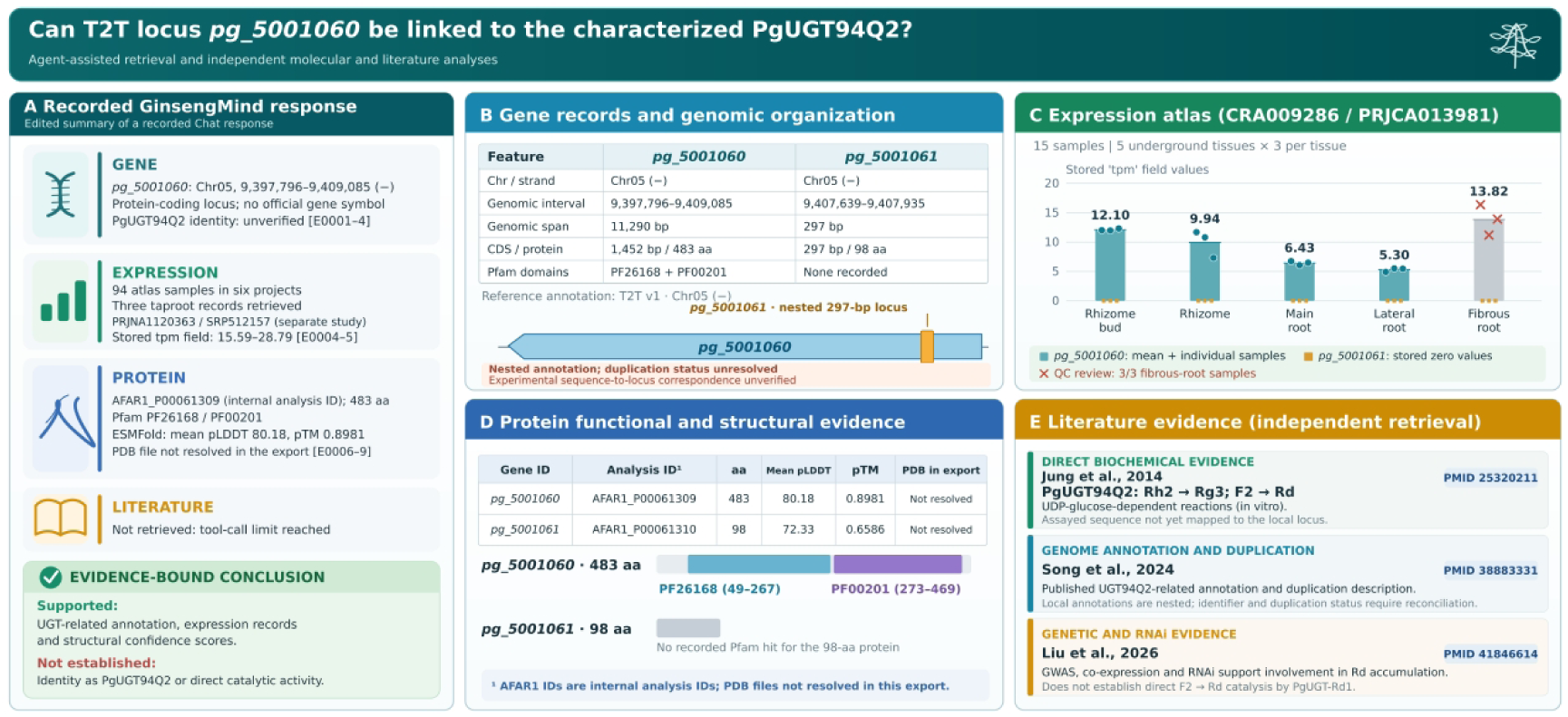
Evidence-grounded analysis of a putative UDP-glycosyltransferase locus in *Panax ginseng* using GinsengMind. (A) An integrated response generated by GinsengMind for the candidate gene *pg_5001060*. The agent retrieved genomic annotations, expression records, protein functional annotations, and ESMFold prediction metrics, providing a source-linked summary of the available evidence. The expression query returned three taproot samples from an atlas comprising 94 samples across six projects. Literature retrieval was not executed in this interaction because the tool-call limit was reached. The response distinguished computational annotations from experimentally validated functions and identified the unresolved correspondence between *pg_5001060* and *PgUGT94Q2*. (B) Genomic organization and annotation of *pg_5001060* and *pg_5001061* based on the *P. ginseng* telomere-to-telomere (T2T) reference genome. Both loci are annotated on the negative strand of chromosome 5. The shorter *pg_5001061* locus (9,407,639–9,407,935 bp) lies within the genomic interval of *pg_5001060* (9,397,796–9,409,085 bp). This overlapping arrangement requires further reconciliation with the tandem-duplication relationship described in the published genome study. (C) Expression profiles of the two candidate genes across five underground tissues using 15 RNA-seq samples from study CRA009286 (BioProject PRJCA013981). Bars represent the mean of three samples per tissue, and individual points indicate the corresponding expression records. The values are derived from the database field labeled tpm. All three fibrous-root samples were flagged for quality-control review. The stored expression values for *pg_5001061* were zero across all 15 samples. Expression-unit provenance, gene-identifier equivalence, and gene-specific read assignment remain to be verified. No statistical significance testing was performed. (D) Protein functional annotations and predicted structural confidence for the two candidate genes. *Pg_5001060* encodes a 483-amino-acid protein annotated with glycosyltransferase-associated Pfam domains PF26168 (residues 49–267) and PF00201 (residues 273–469). The corresponding ESMFold records report mean pLDDT values of 80.18 and 72.33 and pTM scores of 0.8981 and 0.6586 for the proteins associated with *pg_5001060* and *pg_5001061*, respectively. Three-dimensional structures are not displayed because the corresponding PDB files were unavailable in the local export. (E) Literature evidence independently retrieved from the GinsengMind literature database. Jung et al. experimentally characterized *PgUGT94Q2* as a glycosyltransferase catalyzing the conversions of ginsenoside Rh2 to Rg3 and F2 to Rd (20). The 2024 T2T genome study provided genomic annotations and described duplication associated with UGT94Q2 (5), whereas Liu et al. reported genetic association, co-expression, and RNA interference evidence linking *PgUGT-Rd_1_* to ginsenoside Rd accumulation(21). These studies represent distinct levels of functional evidence and were retrieved independently of the Chat interaction shown in (A). The correspondence between the experimentally characterized *PgUGT94Q2* sequence (GenBank accession JX898530) and the reference locus *pg_5001060* remains unverified. Accordingly, the reported catalytic activity of *PgUGT94Q2* cannot yet be directly assigned to this genomic locus.

Genomic examination placed *pg_5001060* on chromosome 5 at positions 9,397,796–9,409,085 on the negative strand, with a 1,452-bp coding sequence encoding a 483-amino-acid protein (Fig. 3B). A second annotated locus, pg_5001061, was located at positions 9,407,639–9,407,935 on the same strand and encoded a substantially shorter 98-amino-acid protein. Notably, the annotated interval of pg_5001061 was entirely contained within that of *pg_5001060*, indicating overlapping rather than two clearly separated genomic loci in the exported reference annotation. Although the published T2T genome study associated UGT94Q2-related annotations with gene duplication, the observed genomic organization requires reconciliation with the original annotation and sequence records before the two loci can be interpreted as tandem duplicates. Furthermore, neither an official PgUGT94Q2 gene-symbol mapping nor a verified correspondence between the previously reported experimental sequence (GenBank accession JX898530) and *pg_5001060* was available in the current integrated records.

The expression atlas provided additional information for examining the candidate locus across underground tissues (Fig. 3C). The original Chat interaction returned three taproot expression records from study PRJNA1120363, with values ranging from 15.59 to 28.79 in the stored tpm field. To examine tissue-associated patterns within a single study, we independently retrieved 15 samples from CRA009286/PRJCA013981, representing rhizome buds, rhizomes, main roots, lateral roots, and fibrous roots, with three samples per tissue. For *pg_5001060*, the corresponding mean stored expression values were 12.10, 9.94, 6.43, 5.30, and 13.82, respectively. By contrast, all 15 records for pg_5001061 contained zero values in the stored expression field. These observations indicate differences in the expression records associated with the two annotated loci; however, they do not independently establish biological expression divergence. In particular, all three fibrous-root samples were flagged for quality-control review, the provenance of the stored expression units had not been independently verified, and the correspondence between expression-atlas identifiers and the T2T reference annotation remained unresolved. No statistical significance testing was performed, and expression measurements from the two independent studies were not combined.

Protein-level analysis further connected the reference loci with sequence-derived functional annotations and stored ESMFold prediction metrics (Fig. 3D). The 483-amino-acid protein associated with *pg_5001060* was indexed as AFAR1_P00061309 and contained two Pfam annotations: PF26168, corresponding to a glycosyltransferase N-terminal domain (residues 49–267), and PF00201, associated with UDP-dependent glycosyltransferases (residues 273–469). Its highest-ranking Swiss-Prot similarity match showed 97.3% sequence identity with 91.5% query coverage. The protein had a mean predicted local distance difference test (pLDDT) score of 80.18 and a predicted template modeling (pTM) score of 0.8981. In comparison, the 98-amino-acid protein associated with pg_5001061 (AFAR1_P00061310) had no recorded Pfam domain and showed a mean pLDDT of 72.33 and a pTM of 0.6586. These findings support a glycosyltransferase-related sequence annotation for the longer protein and provide computational structural-confidence information for both records. However, the corresponding PDB files could not be resolved in the case-study export, preventing direct examination of their predicted three-dimensional structures. Neither the functional annotations nor the prediction scores establish substrate specificity or catalytic activity.

Independent literature retrieval identified three studies providing different levels of evidence relevant to this investigation (Fig. 3E). First, Jung et al. experimentally characterized PgUGT94Q2 and demonstrated its UDP-glucose-dependent conversion of ginsenoside Rh_2_ to Rg_3_ and F_2_ to Rd, providing direct biochemical evidence for these reactions(20). Second, the 2024 T2T genome study(5) associated UGT94Q2-related gene annotations with genomic duplication, although the reported interpretation requires reconciliation with the overlapping coordinates observed in the present reference-data export. Third, a more recent study of PgUGT-Rd_1_ combined genome-wide association analysis, co-expression analysis, and RNA interference to support its involvement in ginsenoside Rd accumulation(21). These genetic and expression-based findings provide evidence of a biological association but do not independently demonstrate direct F_2_-to-Rd catalytic activity by PgUGT-Rd_1_. Importantly, the experimentally characterized PgUGT94Q2 sequence has not yet been conclusively matched to the local *pg_5001060* record; accordingly, its reported enzymatic activities cannot be assigned directly to this reference locus on the basis of the available evidence.

Collectively, the PgUGT94Q2 case illustrates the feasibility of combining GinsengMind-assisted retrieval with independent molecular and literature analyses to investigate a candidate biosynthetic gene. The combined analysis identified unresolved relationships concerning historical sequence-to-genome mapping, overlapping gene annotations, expression-data provenance, and experimentally established enzyme functions. These results highlight the complementary nature of the retrieved molecular evidence and identify specific questions requiring further experimental and computational investigation.

## Discussion

The rapid accumulation of biological data and recent advances in artificial intelligence offer new opportunities to transform how researchers access, integrate, and interpret scientific knowledge. For medicinal plants, however, the effective utilization of molecular resources remains challenging because relevant information is frequently distributed across genomic databases, functional annotation resources, transcriptomic datasets, and experimental literature. Existing resources, such as the Ginseng Genome Database, PanaxGDB, and GinDB-AI, have substantially advanced genomic and chemical investigations of ginseng and related species(11–13). GinsengMind complements these platforms by integrating species-specific molecular knowledge with an evidence-grounded AI research agent, thereby extending conventional database exploration toward question-driven scientific investigation. A distinctive feature of GinsengMind is the combination of a systematically organized molecular knowledge foundation, specialized biological retrieval tools, and LLM-guided evidence interpretation within a unified framework. Rather than treating conversational AI as an auxiliary interface for database retrieval, GinsengMind uses natural-language scientific questions as the starting point for exploring interconnected molecular information. This design provides an approach for transforming biological data resources into interactive research environments in which researchers can investigate molecular relationships and evaluate supporting evidence without relying exclusively on predefined search procedures.

The integration of complementary molecular evidence is particularly important for investigating gene function and specialized metabolism in medicinal plants. Although sequence similarity, orthology-based annotation, and conserved protein domains provide valuable information for functional prediction, the identification of biosynthetic enzymes frequently requires consideration of additional biological and structural evidence. Transcriptomic profiles provide information on the biological contexts in which candidate genes are expressed, whereas predicted protein structures offer opportunities to investigate protein architecture and structural features potentially associated with molecular function. The integration of sequence-derived functional annotations with large-scale protein structure predictions represents an important feature of GinsengMind, enabling researchers to explore complementary protein characteristics within consistent molecular records. For functionally diverse enzyme families involved in specialized metabolism, such as cytochrome P450 monooxygenases and UDP-dependent glycosyltransferases, this integration may facilitate candidate prioritization and provide a starting point for investigating substrate recognition, catalytic diversity, and functional evolution(22, 23). Importantly, the value of integrating multiple evidence sources does not arise simply from accumulating more information, but from establishing traceable relationships among observations while preserving their respective biological meanings and levels of certainty. Combining these molecular features with published experimental findings may therefore support more informed selection of candidate biosynthetic genes and guide subsequent functional characterization.

The GinsengBench evaluation provides quantitative insights into the contribution of domain-specific evidence and autonomous retrieval to scientific question answering. Across the ten core benchmark questions, GinsengMind achieved a mean weighted composite score of 96.18, compared with 23.24 for GLM-5.3 without access to the specialized database or retrieval tools. The substantial improvements in factual accuracy, completeness, and traceability indicate that access to structured molecular information is particularly important for answering species-specific questions that require precise identifiers, quantitative records, and verifiable supporting evidence. The comparison with nine general-purpose LLMs further suggests that broad pretrained knowledge alone may be insufficient for highly specialized scientific tasks. Of particular interest, GLM-5.3 supplied with predefined question-specific evidence packages achieved a mean score of 96.88 across five shared questions, close to the corresponding GinsengMind score of 96.56. Although this comparison does not establish statistical equivalence, it highlights the importance of evidence availability in determining the quality of scientific responses. The value of GinsengMind therefore lies not only in generating accurate scientific answers but also in autonomously acquiring and integrating relevant domain-specific evidence without requiring predefined evidence packages for individual questions. These findings suggest that the development of specialized scientific AI agents should place substantial emphasis on the quality, organization, accessibility, and traceability of domain knowledge, rather than relying exclusively on improvements in the capabilities of underlying language models. In this context, structured retrieval-augmented generation provides a practical approach for connecting general-purpose language models with specialized scientific resources and supporting more reliable biological interpretation.

An important objective of domain-specialized AI research agents is not merely to answer existing scientific questions, but also to assist researchers in identifying knowledge gaps and determining priorities for further investigation. As illustrated by the PgUGT94Q2 case, examining complementary molecular evidence can reveal unresolved relationships that may remain difficult to recognize when information is accessed through isolated queries. Such challenges are particularly relevant to medicinal plant research, where historical biochemical findings, continuously updated genome annotations, and computational predictions may differ in their reference frameworks and levels of certainty. The identification of inconsistencies or incomplete evidence should therefore be regarded not simply as a limitation of information retrieval, but also as an opportunity to refine scientific questions and guide subsequent investigations. For example, unresolved relationships between experimentally characterized enzymes and reference genomic loci can motivate targeted sequence verification, reassessment of gene annotations, and additional biochemical characterization. Although the present case combined Agent-based retrieval with independent data and literature examination, it illustrates how GinsengMind can contribute to a broader evidence-guided research workflow. This approach also highlights the potential of human–AI collaboration in scientific investigation, in which AI agents facilitate evidence acquisition, organization, and interpretation, while researchers retain responsibility for critical scientific judgment, experimental design, and hypothesis validation. Rather than replacing experimental research, such systems may help direct experimental efforts toward more informative biological questions and promising candidate genes.

Beyond ginseng molecular research, the GinsengMind framework has potential implications for medicinal plant specialized metabolism, natural product biosynthesis, and synthetic biology. Despite substantial advances in medicinal plant genomics, the biosynthetic pathways of many pharmacologically important natural products remain incompletely characterized. Identifying the enzymes responsible for their structural diversification often involves extensive searches across heterogeneous molecular datasets, followed by multiple rounds of candidate selection and experimental validation. By connecting functional annotations, expression profiles, structural predictions, and existing biochemical knowledge, domain-specialized AI agents may help streamline these research processes and facilitate the discovery of previously unrecognized biological associations. Integrated molecular evidence may also inform enzyme selection, structure-guided protein engineering, and heterologous biosynthetic pathway reconstruction, thereby contributing to the development of alternative production strategies for valuable natural products. Given its substantial medicinal, scientific, and economic importance, *P. ginseng* provides a representative system for exploring the application of AI agents to medicinal plant research. The modular separation of domain-specific knowledge resources, specialized retrieval tools, and language model interfaces in GinsengMind also offers a potentially adaptable framework for other medicinal plants. More broadly, this approach may inform the development of specialized AI research agents in other scientific disciplines where the interpretation of heterogeneous data, integration of complementary evidence, and formulation of testable hypotheses are central to scientific discovery.

Several limitations of the present study should be acknowledged. First, the reliability of evidence-grounded scientific responses depends on the completeness, quality, and consistency of the underlying molecular resources. Ambiguous gene–transcript–protein relationships, differences among transcriptomic datasets, incomplete structural coverage, and unresolved relationships between historical experimental sequences and reference genome annotations may affect biological interpretation. Continuous literature maintenance and further integration of curated chemical constituents, experimentally characterized biosynthetic reactions, and regulatory information will be important for expanding the scientific scope of GinsengMind. Second, although structured retrieval and evidence management can reduce reliance on unsupported model-generated information, they cannot completely eliminate inaccurate interpretations, incomplete retrieval, or inappropriate biological inference. The current tool-call limits may also restrict the execution of complex, multistep research tasks. Third, although GinsengBench contains 50 curated questions, the present expert evaluation was conducted using ten core questions, with cross-model and fixed-evidence comparisons restricted to five shared questions. Larger independent benchmarks involving diverse research tasks, additional experts, and standardized model configurations will be necessary to establish the robustness and generalizability of the observed performance. Future evaluations should also examine research efficiency, candidate-gene prioritization, and the quality of AI-assisted hypothesis generation under realistic research conditions. Continued improvements in knowledge integration, evidence validation, and autonomous tool orchestration may further advance GinsengMind from a specialized molecular knowledge platform toward an AI-assisted research environment that supports hypothesis-driven investigations. Ultimately, the development of domain-specialized AI research agents provides an opportunity to connect the expanding landscape of scientific data with the experimental processes through which biological knowledge is generated.

## Materials and Methods

### Reference genome and molecular data integration

The molecular resources of GinsengMind were constructed primarily from the published telomere-to-telomere (T2T) reference genome of *Panax ginseng*(5). Gene annotations, transcript sequences, coding sequences, and predicted protein sequences were obtained from the reference genome dataset. Original reference identifiers were retained to maintain consistency with the source annotation. The reference genome assembly and associated annotations were obtained from the publicly available dataset (accession GWHERKY00000000; BioProject PRJCA022032).

Gene, transcript, and protein records were integrated into a relational database. Relationships among these records were established using reference identifiers and sequence compatibility. Ambiguous or unresolved associations were retained separately rather than assigned arbitrarily. Protein sequences containing internal termination symbols were excluded from downstream functional annotation and structure prediction. This procedure established a consistent molecular reference for subsequent data integration and retrieval.

### Protein functional annotation and structure prediction

Protein functional annotations were generated using three complementary approaches. Sequence similarity searches were performed against UniProtKB/Swiss-Prot using DIAMOND, orthology-based annotations were obtained using eggNOG-mapper, and conserved protein families and domains were identified using InterProScan. Annotation records were integrated through standardized protein identifiers while retaining the original database accessions and annotation sources.

Three-dimensional protein structures were predicted using ESMFold with a locally deployed pretrained model. Proteins were processed in length-stratified batches to accommodate computational requirements. The completed prediction set covered proteins up to 800 amino acids in length. Structural models were stored in PDB format, and prediction confidence was recorded using mean pLDDT and pTM scores. Predicted structures and functional annotations were linked to their corresponding protein records to support integrated exploration of sequence, function, and structural information.

### Expression atlas and literature integration

Publicly available *P. ginseng* RNA-seq datasets were organized into a unified expression atlas. Gene-level expression measurements were linked to sample metadata, including study accession, tissue type, experimental conditions, and quality-control status. Dataset provenance was retained to distinguish expression profiles originating from independent studies and to avoid unsupported cross-study comparisons.

A literature resource was established to support retrieval of published studies relevant to ginseng molecular biology and specialized metabolism. Bibliographic records were organized using article metadata and persistent identifiers, including PubMed identifiers (PMIDs) and digital object identifiers (DOIs), where available. A PubMed-based incremental synchronization workflow was developed to support daily literature updates, including the identification of new and revised records, deduplication, and the handling of retracted publications.

### GinsengMind architecture and evidence-grounded AI agent

GinsengMind was implemented using a SQLite-based molecular database and a Streamlit web interface, integrating conventional molecular data retrieval with an LLM-powered conversational research agent. The designated web address of the platform is https://ginsengai.nrc.ac.cn. The system adopts a modular architecture comprising an agent controller, an LLM provider interface, a tool registry, and an evidence management component. The Zhipu GLM-5.3 large language model was used to interpret natural-language queries, coordinate specialized research tools, and generate evidence-grounded scientific responses.

The agent follows a structured retrieval-augmented generation (RAG) workflow in which natural-language questions are interpreted and translated into calls to predefined research tools. Six tools were implemented: search_gene, search_expression, search_protein, search_literature, search_gene_lookup, and search_sequence. These tools provide access to gene records, expression measurements, protein annotations and structural information, scientific literature, gene identifier retrieval, and sequence similarity searches, respectively.

Tool execution is controlled through predefined schemas and authorized retrieval operations. The tool registry validates input parameters and manages access to the corresponding molecular resources. Retrieved records are returned with source identifiers and supporting metadata and subsequently managed by the evidence management component before being supplied to the LLM for response generation. This workflow enables the agent to distinguish database annotations and computational predictions from experimentally supported findings, identify missing evidence, and avoid unsupported functional assignments. Per-query tool and model invocation limits were applied to regulate execution and computational resource consumption.

### GinsengBench design and evaluation

GinsengBench was constructed using 50 curated questions covering gene information, protein annotation and structure, gene expression, literature retrieval, cross-module knowledge integration, and evidence boundaries. Reference answers and permissible inference boundaries were established based on database records and verified tool outputs. Ten core questions spanning all six categories were selected to compare GinsengMind with GLM-5.3 without database access. Five of these questions were additionally used to evaluate nine general-purpose large language models accessed through OpenRouter and five fixed-evidence model configurations. In the fixed-evidence experiments, identical question-specific evidence packages were provided to each model without access to retrieval tools.

A total of 90 responses were anonymized and independently evaluated by three experts against the reference facts and inference boundaries. Each response was scored from 0 to 100 in four dimensions: factual accuracy, completeness, relevance, and traceability, with weights of 40%, 20%, 20%, and 20%, respectively. Scores were averaged across the three experts. Inter-rater agreement was assessed using two-way mixed-effects, absolute-agreement intraclass correlation coefficients (ICC(A,1) and ICC(A,3)). Paired differences were calculated at the question level, and 95% confidence intervals were estimated using 10,000 question-cluster bootstrap resamples. Leave-one-rater-out analysis and an exact paired sign test were performed as supplementary robustness assessments.

### *PgUGT94Q2* case study and evidence assessment

*PgUGT94Q2*, an experimentally characterized UDP-dependent glycosyltransferase involved in ginsenoside biosynthesis, was selected as a representative case to examine the application of GinsengMind to medicinal plant specialized metabolism. The analysis focused on the relationship between the previously characterized enzyme and candidate loci in the *P. ginseng* T2T reference genome.

A genuine GinsengMind Chat interaction was used to retrieve gene annotations, expression records, protein functional information, and structural prediction metrics associated with the candidate locus. Additional expression records, genomic annotations, and relevant publications were independently examined to extend the analysis beyond the information retrieved during the original interaction.

For the expression analysis, 15 RNA-seq samples from five underground tissue types were retrieved from study CRA009286 (BioProject PRJCA013981), with three samples per tissue. Expression values were extracted from the existing atlas and summarized as arithmetic means for descriptive comparison. Sample-level quality-control flags were retained, and no statistical significance testing was performed.

Protein-level evidence was examined using the corresponding internal protein records, including sequence length, functional annotations, conserved domains, and ESMFold confidence metrics. Literature evidence was independently reviewed to distinguish experimentally demonstrated enzyme activities from genome-based annotations, expression associations, and computational predictions.

The published PgUGT94Q2 sequence (GenBank accession JX898530) was considered in evaluating its potential correspondence to the candidate T2T loci. Because a definitive sequence-to-locus assignment had not been established, the candidate loci were not treated as confirmed genomic representations of PgUGT94Q2. All conclusions were interpreted according to the corresponding level of experimental or computational evidence.

## Data and code availability

The genomic and transcriptomic datasets used in this study were obtained from publicly available repositories. The integrated molecular resources are accessible through the GinsengMind platform (https://ginsengai.nrc.ac.cn). Data and materials supporting the GinsengBench evaluation and case study are available from the corresponding author upon reasonable request. Additional information regarding the implementation and availability of the platform and associated code can also be obtained from the corresponding author.

## Acknowledgements

We thank Zhixian Jing and Tielin Wang from the National Resource Center for Chinese Materia Medica, China Academy of Chinese Medical Sciences, for their valuable support and assistance.

## CRediT authorship contribution statement

**Yuru Tong:** Conceptualization, Methodology, Writing – original draft, Software, Resources, Funding acquisition; **Rong Zhang:** Conceptualization, Methodology; **Chuanzhi Kang:** Resources, Funding acquisition; **Zeqian Chang:** Resources; **Kang Chen:** Conceptualization, Methodology, Formal analysis, Investigation, Writing – original draft, Writing – Review & Editing, Software, Visualization

